# scAmpi - A versatile pipeline for single-cell RNA-seq analysis from basics to clinics

**DOI:** 10.1101/2021.03.25.437054

**Authors:** Anne Bertolini, Michael Prummer, Mustafa Anil Tuncel, Ulrike Menzel, María Lourdes Rosano-González, Jack Kuipers, Daniel Johannes Stekhoven, Tumor Profiler consortium, Niko Beerenwinkel, Franziska Singer

## Abstract

Single-cell RNA sequencing (scRNA-seq) has emerged as a powerful technique to decipher tissue composition at the single-cell level and to inform on disease mechanisms, tumor heterogeneity, and the state of the immune microenvironment. Although multiple methods for the computational analysis of scRNA-seq data exist, their application in a clinical setting demands standardized and reproducible workflows, targeted to extract, condense, and display the clinically relevant information. To this end, we designed scAmpi (**S**ingle **C**ell **A**nalysis **m**RNA **pi**peline), a workflow that facilitates scRNA-seq analysis from raw read processing to informing on sample composition, clinically relevant gene and pathway alterations, and *in silico* identification of personalized candidate drug treatments. We demonstrate the value of this workflow for clinical decision making in a molecular tumor board as part of a clinical study.

## Introduction

In recent years, single-cell RNA sequencing (scRNA-seq) emerged as a high-throughput technology for uncovering gene expression at the single-cell level, which provides unprecedented insights into, e.g., cell differentiation, the immune compartment, and tumor heterogeneity [1, 2]. Initially used to characterize PBMCs or differentiating stem cells, an increasing number of studies exploit scRNA-seq to investigate clinical samples such as tumor tissues [3, 4]. There are multiple software suites available with extensive functionality for general scRNA-seq analysis, including the widely-used tools SEURAT [5] and ScanPy [6]. However, they have some disadvantages: First, for non-bioinformaticians the usage can be difficult because setting all parameters and applying the different steps requires at least basic R or Python programming knowledge. Second, to the best of our knowledge, no software is available that facilitates *in-silico* drug candidate identifications based on single-cell data. Finally, existing software suites are not designed to manage large-scale data analysis in a highly reproducible, transparent, and auditable way, including error tracking and process documentation, and thus are not suitable to be employed for routine clinical use [19, 20]. We therefore developed scAmpi, an end-to-end turn-key pipeline for scRNA-seq analysis from raw read processing to informing on sample composition, gene expression, and potential drug candidates. Utilizing the Snakemake workflow management system [7], scAmpi is easy to use and offers a high degree of flexibility in the choice of methods, while it can be employed in a highly standardized and reproducible fashion. This has led to the successful implementation of scAmpi for processing scRNA-seq data in the ongoing Tumor Profiler clinical study [8, 21].

## Design and Implementation

In the following, we describe how scAmpi can be used for analyzing tumor scRNA-seq data from the 10x Genomics platform (Fig 1). All workflow steps, such as e.g., choice of the read mapper or clustering method, can be replaced with little effort and the workflow is directly applicable also to other tissue types. A complete analysis of a single sample with approx. 4,000 cells and 50,000 reads per cell takes four to eight hours, depending on the available compute resources. The pipeline can easily scale to the parallel analysis of large cohorts of hundreds of samples, where each sample is processed independently in a single-sample analysis fashion. To ensure a thoroughly documented analysis, each workflow step is tracked with log files describing command, input, output, and resource requirements, as well as error documentation.

**Figure 1:**
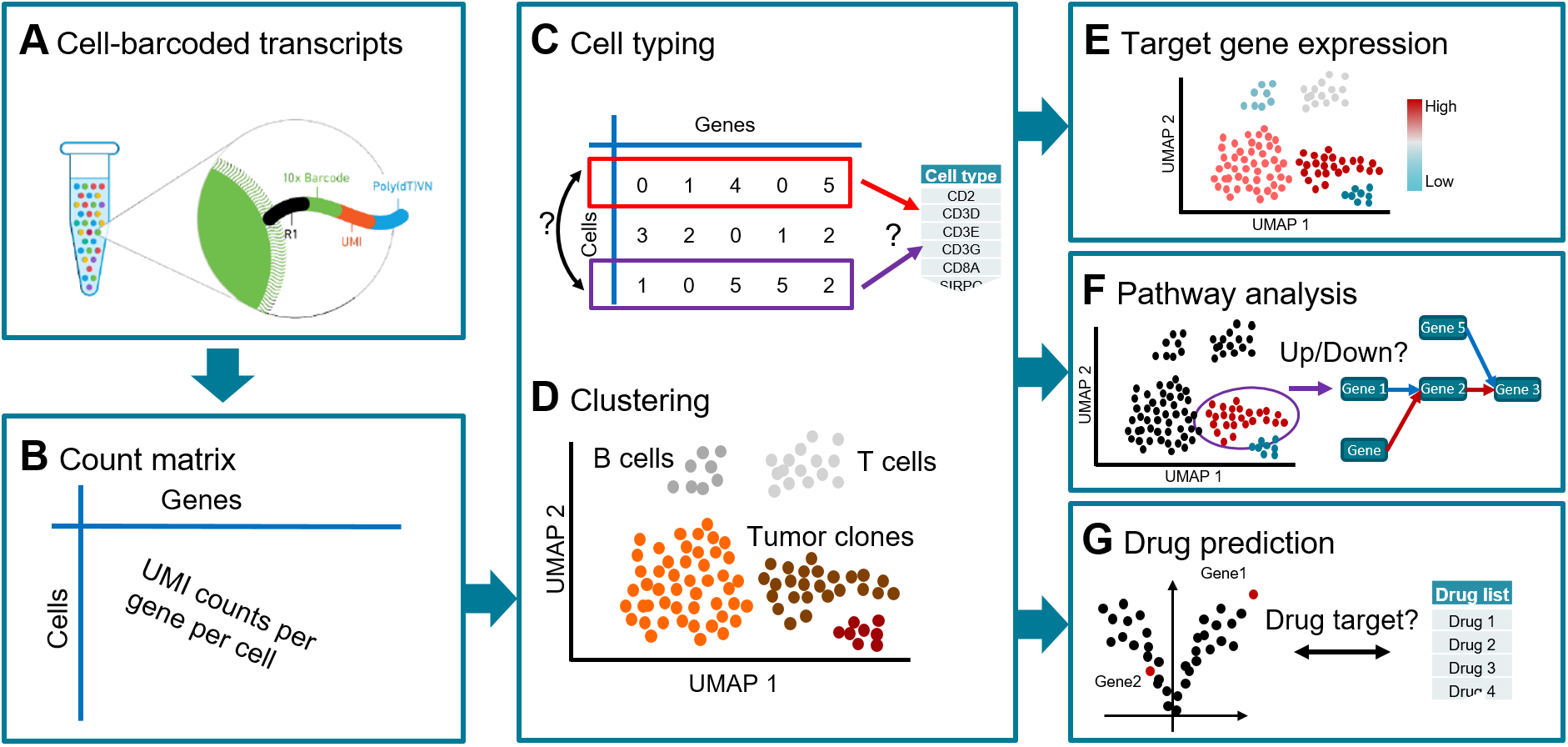
Overview of the workflow implemented in scAmpi, showing a tumor sample analysis as an example. Starting from droplet-based 10x Genomics raw data (A), genome-wide read counts for each cell are generated (B). This gene-by-cell count matrix is the basis for cell type prediction (C) and unsupervised clustering (D) to determine the cell type composition and tumor heterogeneity. Subsequent steps include gene expression (E) and gene set (F) analysis, and drug candidate identification (G).

### Read data processing and normalization

Using the Cellranger software, reads are assigned to their respective cells based on the 10x Genomics barcodes and simultaneously mapped to the reference genome to infer read counts per gene per cell. Subsequently, several filters are applied to remove contaminants, cell fragments, or dying cells. Per default, all non-protein-coding genes and genes coding for ribosomal proteins are removed. All cells exceeding a specific threshold in the number of reads mapping to mitochondrial genes are discarded, because they are likely broken cells [9]. This threshold can be either user-specified or estimated from the data. Further, all cells with too few expressed genes are discarded in order to remove low-quality cells or presumably empty droplets. The remaining counts are normalized for cell-cycle effect and library size using sctransform [10], which yields Pearson residuals as well as corrected counts per gene per cell for the subsequent analysis steps. Several types of plots, including scatter plots showing cells that are filtered out and box plots showing the most highly expressed genes in the sample, are provided to support quality control.

### Sample composition

The two key analyses to inform on sample composition are cell type identification (Fig 1C) and unsupervised clustering (Fig 1D). Per default, clustering is performed using Phenograph [11]. The clustering compares expression profiles across cells and yields groups of highly similar cells. Per default, a minimum of 20 cells per group are required, in order to reach a group size suitable for subsequent differential gene expression analysis.

In contrast, automated cell type classification is applied to each cell individually (manuscript in preparation). Briefly, the expression profile of each cell is compared to *a priori* defined lists of cell type marker genes. Each cell type is represented by a list of genes that are known to be specific for and highly expressed in this cell type. The set of cell types used for classification is expected to reflect the cell types present in the analyzed tissue. The method accommodates for uncertainty in the typing as well as unknown cell types. If the expression profile of a cell does not reach a specified similarity threshold, it is labeled as ‘unknown’. If a cell matches two or more cell types with high similarity (i.e., the best and second-best similarity scores are too similar), it is typed as ‘uncertain’. Cell type lists can be derived from literature. For example, for melanoma biopsies, we based our typing on the markers published by Tirosh et al. [3]. Using a cell type list that was derived from data of another tissue is also possible, but should be done with care due to tissue-dependent expression differences. The cell type analysis works in a two-step hierarchical fashion. In the first iteration the major cell type populations are identified, e.g., tumor cells are distinguished from T cells. In a second step, all cells belonging to a particular major cell type can be re-classified into subtypes. For instance, T cells can be sub-classified (among others) into gamma delta, memory resting, or regulatory T cells. scAmpi already offers predefined cell type lists for melanoma, AML, ovarian cancer, and PBMCs, but user-specified marker lists can be easily added.

The results of the sample composition analysis (unsupervised clustering and cell typing) are visualized in a low-dimensional representation using, e.g., Uniform Manifold Approximation and Projection (UMAP) [18].

### Differential gene expression

Detecting differential gene expression (DE) is a major aspect of standard mRNA sequencing experiments. Here, we perform two main comparisons for scRNA-seq data using multiple linear regression: First, provided multiple tumor clusters are found, a DE analysis is performed that compares the expression phenotypes of the different tumor clusters and informs on the tumor heterogeneity. Second, provided malignant (tumor) cells as well as non-malignant cells are found, a DE analysis is performed that identifies genes with different expression levels in each tumor cluster compared to all non-malignant cells. Non-malignant cells can be any cell type present in the tissue, such as, immune cells, endothelial, or epithelial cells.

### Gene expression and pathway analysis

The user can provide grouped lists of priority genes and pathways to be visualized (Fig 1E). Gene expression is visualized for each cell in a color-coded UMAP together with a violin plot that shows the expression distribution per cluster, separately for each group of genes. Further, for each cluster, various gene expression summary statistics are provided, such as the gene expression rank, the average expression, and the proportion of cells with non-zero expression.

Pathway analysis is performed in two independent approaches (Fig 1F). Based on the DE genes, a competitive gene set analysis is performed using the camera function from the limma R package [12]. Here, we output all pathways with an FDR below a user-defined cut-off that are up-regulated, down-regulated, or are categorized as mixed if both over- and under-expressed genes were identified in the respective pathway. Gene set enrichment based on DE analysis is very common, but has certain drawbacks for single-cell data, as these experiments often lack a proper reference, which can bias the pathway enrichment. Thus, we also perform a GSVA-based pathway analysis [13], in which gene sets are ranked relative to each other within each cell independent of all other cells. As this approach is comparing gene sets within a cell, it does not rely on the presence of a reference cell population.

### In-silico drug candidate identification

Initially developed for bulk sequencing data [14], the *in-silico* drug candidate identification framework was refined and adapted to facilitate single-cell and expression data analysis (Fig 1G). For each tumor cell cluster, the differentially expressed genes resulting from the comparison of malignant versus non-malignant cells are used to query DGIdb [15] to obtain potential drug-gene interactions. These drug-gene interactions are undirected in the sense that they do not reveal whether the tumor might be sensitive or resistant to the identified drug. Thus, we further enrich the drug-gene interactions with information from clinicaltrials.gov and CIViC [16]. CIViC is a database of curated drug-gene interaction information providing information on the observed expression type, i.e., over-expression or under-expression. This directed *in-silico* drug candidate identification is also visualized on the sample composition UMAP.

## Results

We showcase the readout and analyses possible with scAmpi for scRNA-seq data from a biopsy of a melanoma patient who was included in the Tumor Profiler clinical study [8]. The full analysis from raw fastq files to *in-silico* drug candidate identification is triggered with only two commands. For details on the default parameter settings, we refer to S1. In the initial mapping step, Cellranger identifies 4193 cells. Subsequent filtering in scAmpi removes 10% (437) of cells due to low quality (Figure 2A). Figures 2B and 2C show examples of QC metrics on the UMAP representation of the cells. After normalization, the cell-cycle phase has no apparent effect on the embedding of the cells anymore. Instead, as shown in Figure 3A, the embedding is cleanly separated by cell type populations.

**Figure 2:**
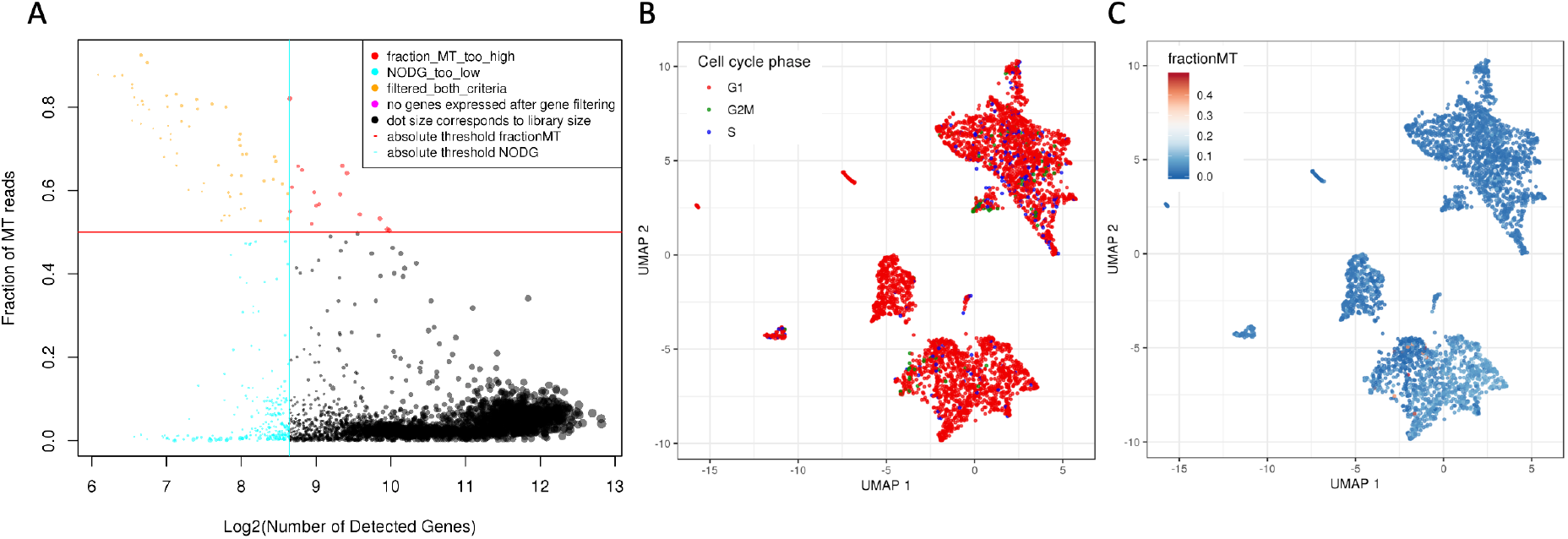
Examples of scAmpi’s basic scRNA-seq quality control plots of a melanoma sample. The scatter plot in (A) shows cells colored by their respective category of applied filters. The vertical and horizontal lines indicate the chosen thresholds applied for the minimum number of genes (x-axis) and maximum fraction of reads mapping to mitochondrial genes per cell (y-axis), respectively. In (B), the UMAP embedding (after normalization) of all cells is shown, with cells colored by estimated cell-cycle phase. In (C), the same UMAP is shown, this time with cells colored by the fraction of reads mapping to mitochondrial genes.

The cell type composition analysis identifies a melanocytic melanoma cell population that constitutes 34% of the sample. The tumor immune microenvironment is very diverse and shows a large group of T cells, mainly sub-classified as memory effector T cells, as well as macrophages, B cells, NK cells, and Endothelial cells (Figure 3A). This finding is in agreement with results from CyTOF experiments also performed on the case study presented in [8]. Further investigation of the immune microenvironment is facilitated by gene expression visualization and a population-based and ranked overview of the average gene expression and number of non-zero cells for each gene. For instance, as shown in Supp. Figure 2, memory effector T cells express PDCD1 (PD-1), an immune checkpoint marker relevant for immunotherapy. Other immune checkpoint markers are also expressed, together with an observed MHC class I expression (HLA-A/B/C) on the tumor cells indicating that T cells would be able to recognize tumor cells. Taken together, this molecular phenotype suggests a potential suitability of anti-PD1 immunotherapy. This finding is also supported by other technologies presented in [8], such as CyTOF and imaging mass cytometry.

**Figure 3:**
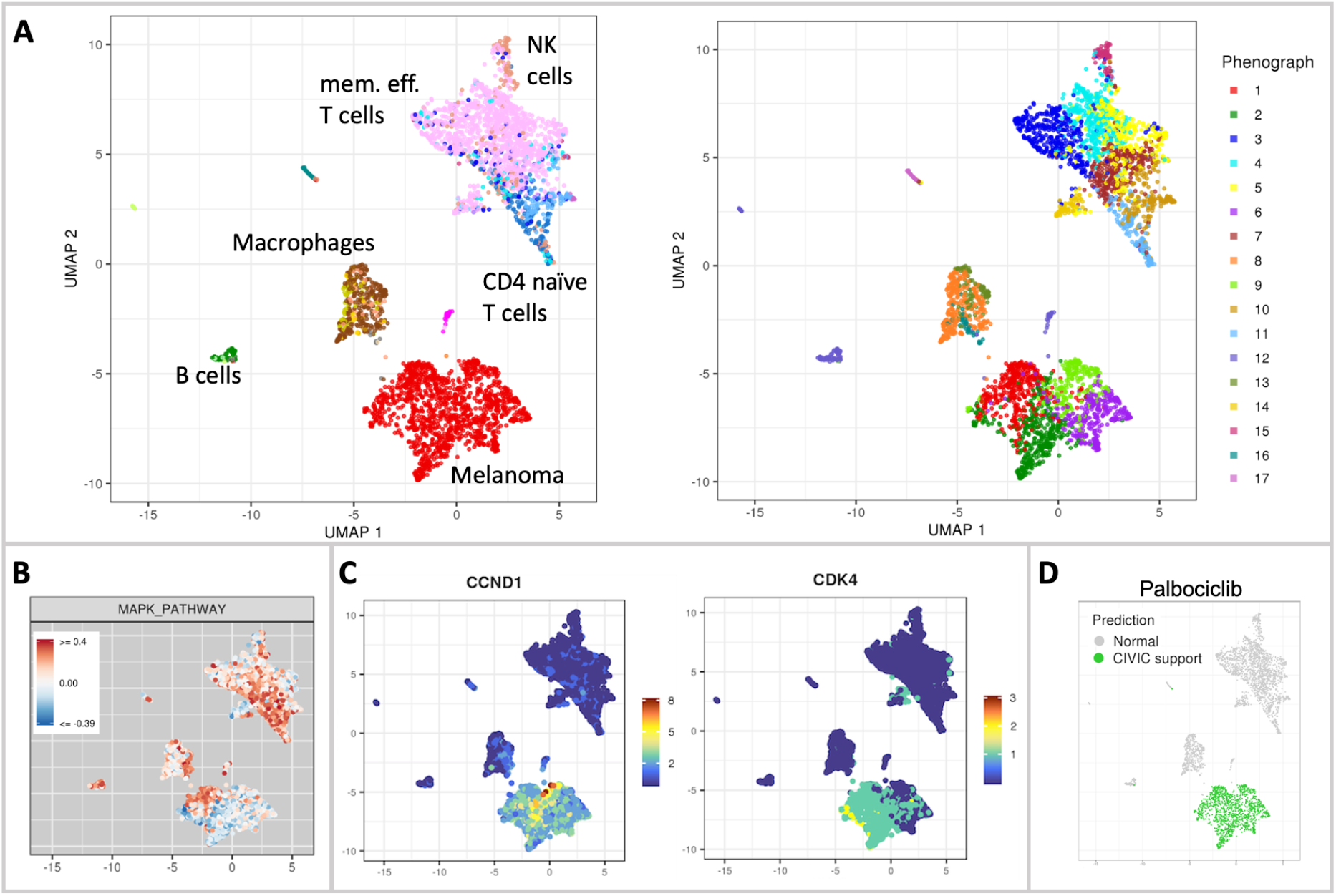
Sample composition and interpretation of a melanoma sample. In (A) the UMAP embedding is colored by cell type label (left) and cluster (right), with major cell type populations highlighted in the figure. For a complete overview of cell types, see Supp. Figure 3. In (B), the enrichment of the MAPK pathway is exemplified. In (C), UMAPs showing the gene expression of CCND1 and CDK4 are shown as selected examples of individual gene expression plots. The UMAP in (D) shows the drug candidate identification result for the drug palbociclib.

Unsupervised clustering (Figure 3A) reveals that the melanoma population groups into four clusters, indicating tumor heterogeneity. scAmpi offers multiple readouts to further investigate this heterogeneity, including, e.g., individual gene expression analysis, gene set enrichment analysis, and differential gene expression comparing the tumor clusters (see S3 for details). As shown in Figure 3B, three of the four tumor clusters display down-regulation of the MAPK pathway (gene set taken from the Hallmark MSigDB [17]), precluding the use of BRAF/MEK inhibitor treatment. In contrast, the *in-silico* drug candidate identification of scAmpi marked the complete tumor population to be potentially sensitive to palbociclib treatment, based on the over-expression of CCND1 and further supported by the expression of CDK4 (Figures 3C and 3D). This finding is observed across other technologies described in [8], such as drug response testing (referred to as Pharmacoscopy).

Taken together, scAmpi provides not only insights into the general sample composition and gene and pathway expression, but also enables downstream data interpretation to support clinical decision making.

## Availability and Future Directions

The source code of scAmpi is available on github at https://github.com/ETH-NEXUS/scAmpi_single_cell_RNA. scAmpi offers comprehensive functionality for the analysis of scRNA-seq data. Key aspects are on the one hand its flexibility and ease of use, which allows the application to various tissues and disease types. On the other hand, it provides a standardized and reproducible workflow that is suited for application in clinical settings and was already utilized in a clinical study [8, 21]. Moreover, scAmpi facilitates *in-silico* drug candidate identification on the single-cell level, thereby directly accounting for disease heterogeneity in the design of optimal drug treatment. Finally, because of the modular Snakemake framework, we foresee a continued extension and refinement of the pipeline and its open source code, also by the single-cell community.

## Acknowledgements

The study described in this paper is the result of a jointly-funded effort between several academic institutions (The University of Zurich, The University of Zurich Hospital, The Swiss Federal Institute of Technology in Zurich, The University of Basel Hospital), as well as F. Hoffmann-La Roche AG.

## Ethics approval

Ethics approval has been granted: BASEC-Nr.2018-02050.

## Authors contributions

AB, MP, and FS designed and implemented the pipeline. MLRG and MAT contributed features to the implementation. UM provided cell-type classification support. AB, MP, JK, DS, NB, and FS wrote and revised the manuscript. All authors have read and approved the manuscript.

