## Supplementary material for "scAmpi - A versatile pipeline for single-cell RNA-seq analysis from basics to clinics": Supp.

**S1: Parameter setting and analysis call**

Basic scRNA:
```
snakemake -s snake_scAmpi_basic_master.snake --configfile
config_MelanomaSample.json
```

Clinical:
```
snakemake -s snake_scAmpi_clinical_master.snake --configfile
config_MelanomaSample.json
```

Analysis-specific parameters and resources required by the pipeline can be provided in a configuration file in json format. An example file listing the default settings is provided on the git repository.

Specific parameters for the melanoma showcase example:
Normalization:
MT-fraction threshold = 0.5
Number of genes = 400

Clustering:
Number of neighbors = 30
Minimum number of cells per cluster = 20
Number of highly variable genes = 2000

Clinical part:
clinicaltrials.gov key words: “solid tumor, melanoma”

### S2: Cell types and immune gene expression

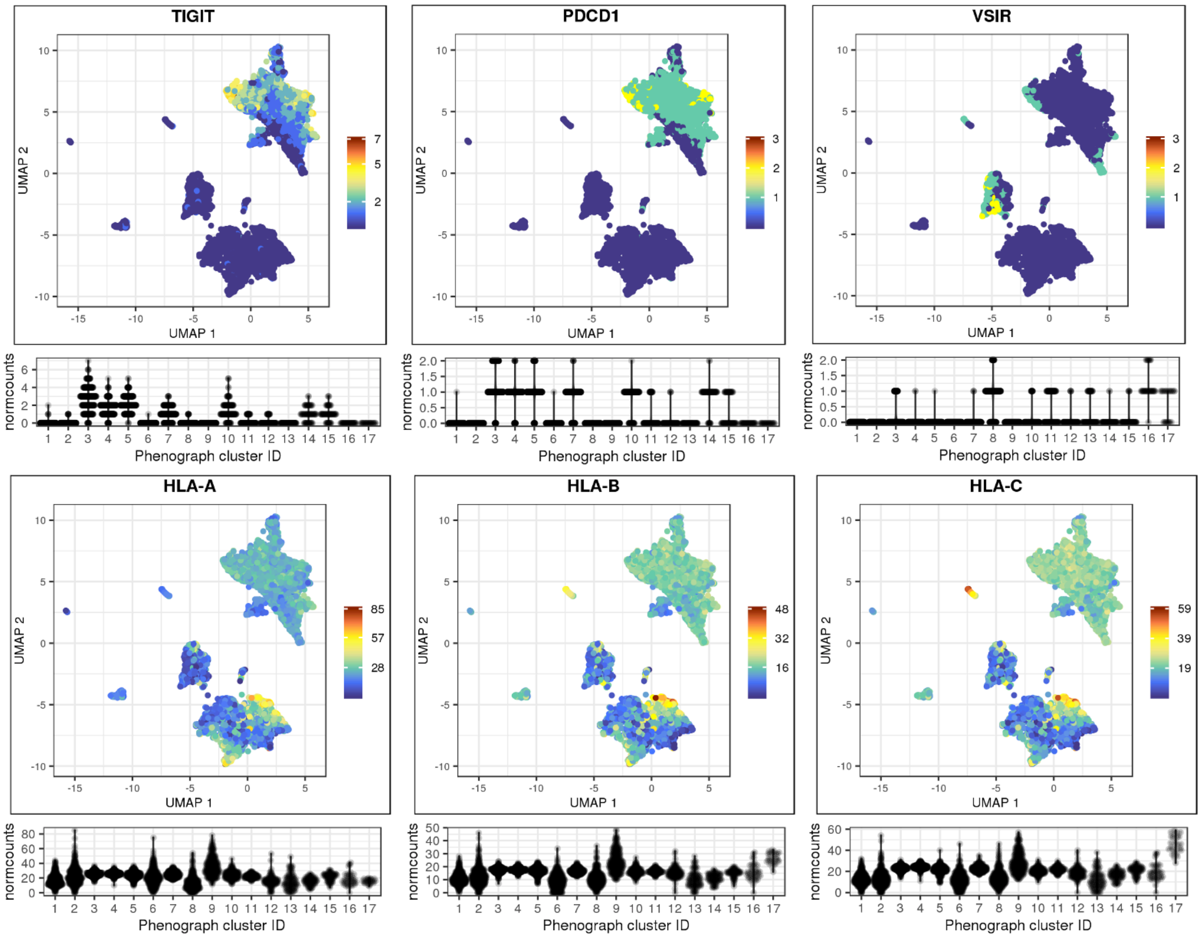


*Supplementary Figure 2: Single cell UMAPs and violin plots showing the normalized expression counts of immunotherapy-relevant marker genes (gene to protein names: PDCD1 = PD-1; VSIR = VISTA).*

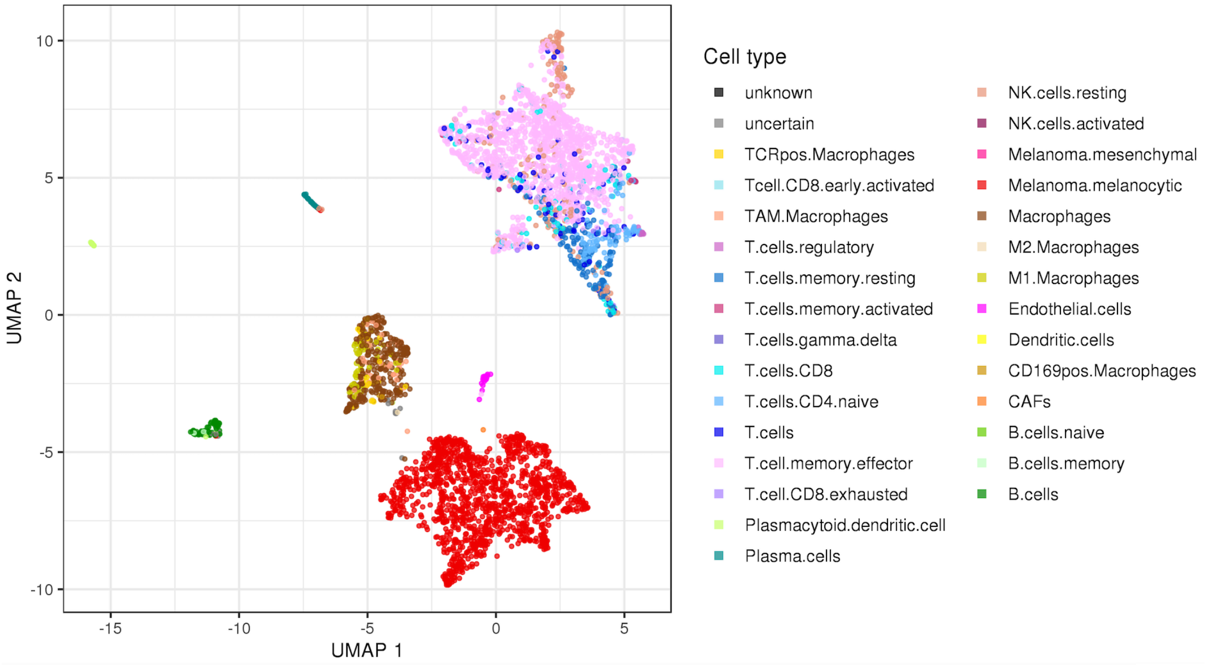


*Supplementary Figure 3: Complete overview of the cell type composition of a melanoma biopsy sample.*

### S3: Tumor heterogeneity

Supplementary Figures 4 and 5 show selected examples of the gene set enrichment analysis based on differentially expressed genes and GSVA, respectively. The heat map illustrates that e.g. Interferon alpha/gamma response are up-regulated in cluster 9 but down-regulated in cluster 6. In combination with the GSVA score-colored UMAP it becomes apparent that the tumor is generally down-regulated for interferon alpha/gamma, with the exception of cluster 9.

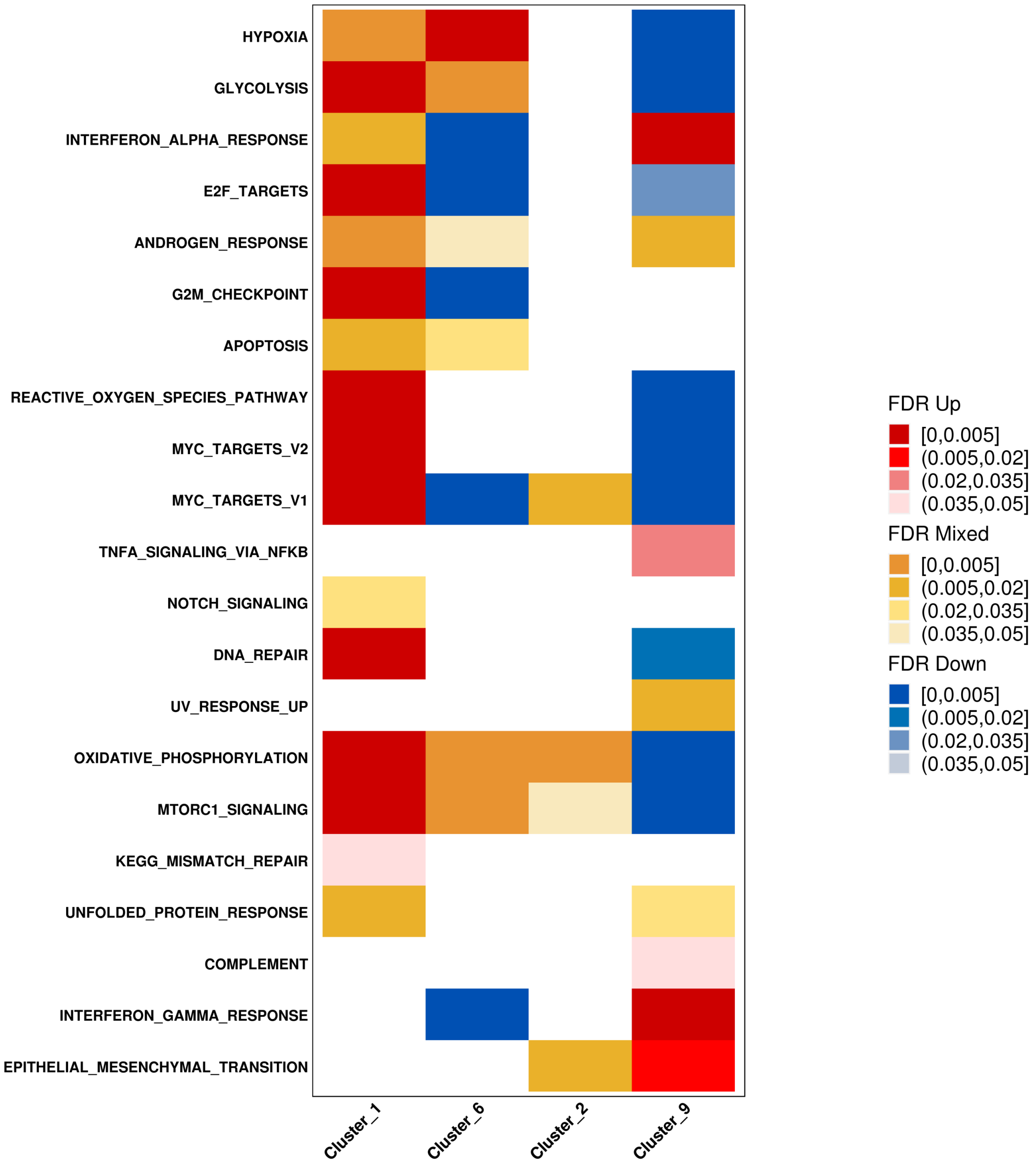


*Supplementary Figure 4: Heat map illustrating the gene set enrichment analysis results based on genes differentially expressed comparing malignant clusters with each other.*

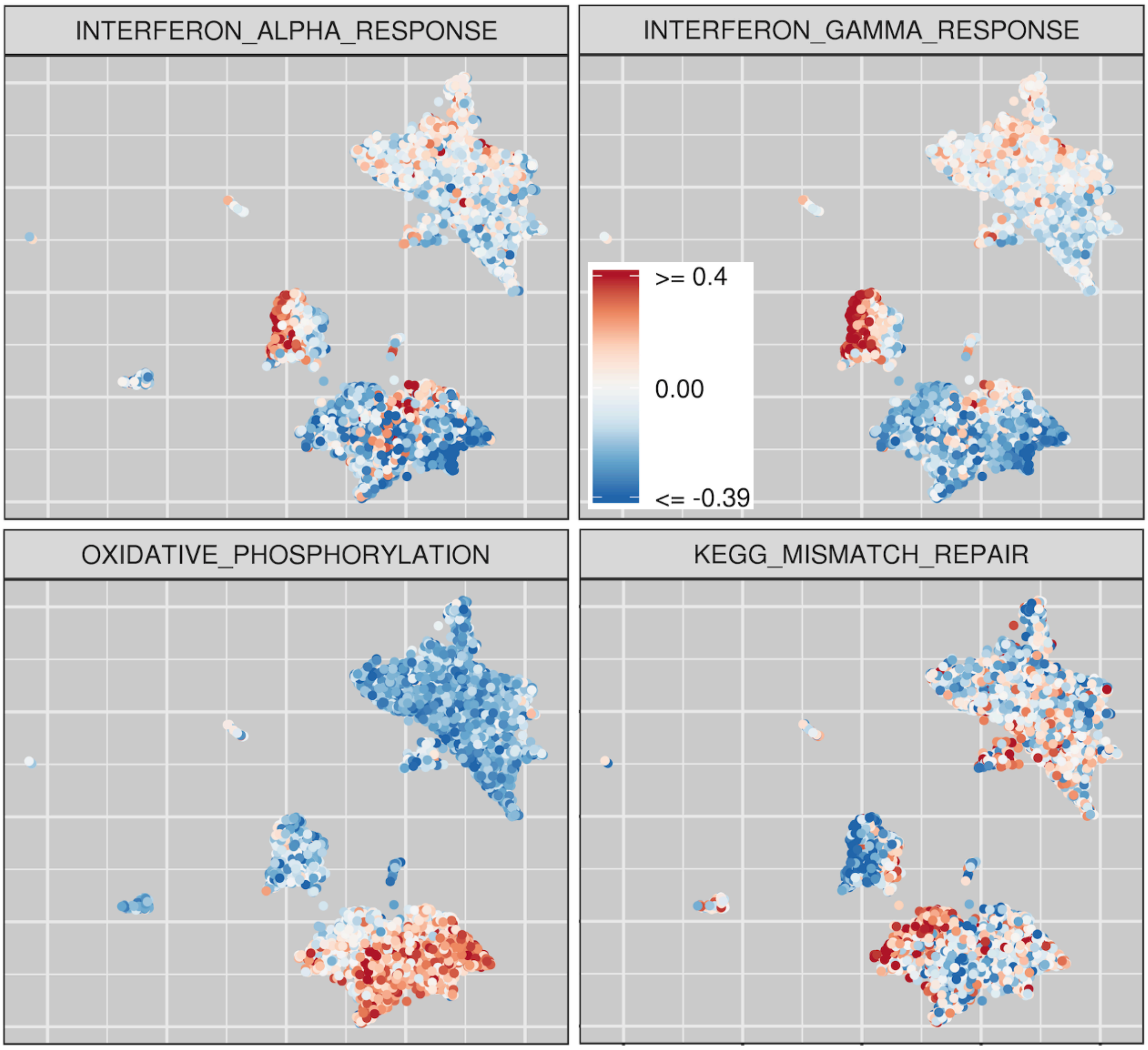


*Supplementary Figure 5: UMAPs that illustrate selected examples of the GSVA based gene set enrichment analysis.*

### TUPRO Consortium

Rudolf Aebersold[2], Melike Ak[27], Faisal S Al-Quaddoomi[9,16], Jonas Albinus[7], Ilaria Alborelli[23], Sonali Andani[6,16,25,30], Per-Olof Attinger[11], Marina Bacac[15], Daniel Baumhoer[23], Beatrice Beck-Schimmer[38], Niko Beerenwinkel[4,16], Christian Beisel[4], Lara Bernasconi[26], Anne Bertolini[9,16], Bernd Bodenmiller[8,34], Ximena Bonilla[6,16,25], Lars Bosshard[9,16], Byron Calgua[23], Ruben Casanova[34], Stéphane Chevrier[34], Natalia Chicherova[9,16], Maya D'Costa[10], Esther Danenberg[36], Natalie Davidson[6,16,25], Monica-Andreea Drăgan[4], Reinhard Dummer[27], Stefanie Engler[34], Martin Erkens[13], Katja Eschbach[4], Cinzia Esposito[36], André Fedier[17], Pedro Ferreira[4], Joanna Ficek[6,16,25], Anja L Frei[30], Bruno Frey[12], Sandra Goetze[7], Linda Grob[9,16], Gabriele Gut[36], Detlef Günther[5], Martina Haberecker[30], Pirmin Haeuptle[1], Viola Heinzelmann-Schwarz[17,22], Sylvia Herter[15], Rene Holtackers[36], Tamara Huesser[15], Anja Irmisch[27], Francis Jacob[17], Andrea Jacobs[34], Tim M Jaeger[11], Katharina Jahn[4], Alva R James[6,16,25], Philip M Jermann[23], André Kahles[6,16,25], Abdullah Kahraman[16,30], Viktor H Koelzer[30], Werner Kuebler[24], Jack Kuipers[4,16], Christian P Kunze[21], Christian Kurzeder[20], Kjong-Van Lehmann[6,16,25], Mitchell Levesque[27], Sebastian Lugert[10], Gerd Maass[12], Markus G Manz[29], Philipp Markolin[6,16,25], Julien Mena[2], Ulrike Menzel[4], Julian M Metzler[28], Nicola Miglino[1], Emanuela S Milani[7], Holger Moch[30], Simone Muenst[23], Riccardo Murri[37], Charlotte KY Ng[23,33], Stefan Nicolet[23], Marta Nowak[30], Patrick GA Pedrioli[3], Lucas Pelkmans[36], Salvatore Piscuoglio[17,23], Michael Prummer[9,16], Mathilde Ritter[17], Christian Rommel[13], María L Rosano-González[9,16], Gunnar Rätsch[3,6,16,25], Natascha Santacroce[4], Jacobo Sarabia del Castillo[36], Ramona Schlenker[14], Petra C Schwalie[13], Severin Schwan[11], Tobias Schär[4], Gabriela Senti[26], Franziska Singer[9,16], Sujana Sivapatham[34], Berend Snijder[2,16], Bettina Sobottka[30], Vipin T Sreedharan[9,16], Stefan Stark[6,16,25], Daniel J Stekhoven[9,16], Alexandre PA Theocharides[29], Tinu M Thomas[6,16,25], Markus Tolnay[23], Vinko Tosevski[15], Nora C Toussaint[9,16], Mustafa A Tuncel[4,16], Marina Tusup[27], Audrey Van Drogen[7], Marcus Vetter[19], Tatjana Vlajnic[23], Sandra Weber[26], Walter P Weber[18], Rebekka Wegmann[2], Michael Weller[32], Fabian Wendt[7], Norbert Wey[30], Andreas Wicki[29,35], Mattheus HE Wildschut[2,29], Bernd Wollscheid[7], Shuqing Yu[9,16], Johanna Ziegler[27], Marc Zimmermann[6,16,25], Martin Zoche[30], Gregor Zuend[31]

[1]Cantonal Hospital Baselland, Medical University Clinic, Rheinstrasse 26, 4410 Liestal, Switzerland, [2]ETH Zurich, Department of Biology, Institute of Molecular Systems Biology, Otto-Stern-Weg 3, 8093 Zurich, Switzerland, [3]ETH Zurich, Department of Biology, Wolfgang-Pauli-Strasse 27, 8093 Zurich, Switzerland, [4]ETH Zurich, Department of Biosystems Science and Engineering, Mattenstrasse 26, 4058 Basel, Switzerland, [5]ETH Zurich, Department of Chemistry and Applied Biosciences, Vladimir-Prelog-Weg 1-5/10, 8093 Zurich, Switzerland, [6]ETH Zurich, Department of Computer Science, Institute of Machine Learning, Universitätstrasse 6, 8092 Zurich, Switzerland, [7]ETH Zurich, Department of Health Sciences and Technology, Otto-Stern-Weg 3, 8093 Zurich, Switzerland, [8]ETH Zurich, Institute of Molecular Health Sciences, Otto-Stern-Weg 7, 8093 Zurich, Switzerland, [9]ETH Zurich, NEXUS Personalized Health Technologies, John-von-Neumann-Weg 9, 8093 Zurich, Switzerland, [10]F. Hoffmann-La Roche Ltd, Grenzacherstrasse 124, 4070 Basel, Switzerland, [11]F. Hoffmann-La Roche Ltd, Grenzacherstrasse 124, 4070 Basel, Switzerland, , [12]Roche Diagnostics GmbH, Nonnenwald 2, 82377 Penzberg, Germany, [13]Roche Pharmaceutical Research and Early Development, Roche Innovation Center Basel, Grenzacherstrasse 124, 4070 Basel, Switzerland, [14]Roche Pharmaceutical Research and Early Development, Roche Innovation Center Munich, Roche Diagnostics GmbH, Nonnenwald 2, 82377 Penzberg, Germany,

[15]Roche Pharmaceutical Research and Early Development, Roche Innovation Center Zurich, Wagistrasse 10, 8952 Schlieren, Switzerland, [16]SIB Swiss Institute of Bioinformatics, Lausanne, Switzerland, [17]University Hospital Basel and University of Basel, Department of Biomedicine, Hebelstrasse 20, 4031 Basel, Switzerland, [18]University Hospital Basel and University of Basel, Department of Surgery, Brustzentrum, Spitalstrasse 21, 4031 Basel, Switzerland, [19]University Hospital Basel, Brustzentrum & Tumorzentrum, Petersgraben 4, 4031 Basel, Switzerland, [20]University Hospital Basel, Brustzentrum, Spitalstrasse 21, 4031 Basel, Switzerland, [21]University Hospital Basel, Department of Information- and Communication Technology, Spitalstrasse 26, 4031 Basel, Switzerland, [22]University Hospital Basel, Gynecological Cancer Center, Spitalstrasse 21, 4031 Basel, Switzerland, [23]University Hospital Basel, Institute of Medical Genetics and Pathology, Schönbeinstrasse 40, 4031 Basel, Switzerland, [24]University Hospital Basel, Spitalstrasse 21/Petersgraben 4, 4031 Basel, Switzerland, [25]University Hospital Zurich, Biomedical Informatics, Schmelzbergstrasse 26, 8006 Zurich, Switzerland, [26]University Hospital Zurich, Clinical Trials Center, Rämistrasse 100, 8091 Zurich, Switzerland, [27]University Hospital Zurich, Department of Dermatology, Gloriastrasse 31, 8091 Zurich, Switzerland, [28]University Hospital Zurich, Department of Gynecology, Frauenklinikstrasse 10, 8091 Zurich, Switzerland, [29]University Hospital Zurich, Department of Medical Oncology and Hematology, Rämistrasse 100, 8091 Zurich, Switzerland, [30]University Hospital Zurich, Department of Pathology and Molecular Pathology, Schmelzbergstrasse 12, 8091 Zurich, Switzerland, [31]University Hospital Zurich, Rämistrasse 100, 8091 Zurich, Switzerland, [32]University Hospital and University of Zurich, Department of Neurology, Frauenklinikstrasse 26, 8091 Zurich, Switzerland, [33]University of Bern, Department of BioMedical Research, Murtenstrasse 35, 3008 Bern, Switzerland, [34]University of Zurich, Department of Quantitative Biomedicine, Winterthurerstrasse 190, 8057 Zurich, Switzerland, [35]University of Zurich, Faculty of Medicine, Zurich, Switzerland, [36]University of Zurich, Institute of Molecular Life Sciences, Winterthurerstrasse 190, 8057 Zurich, Switzerland, [37]University of Zurich, Services and Support for Science IT, Winterthurerstrasse 190, 8057 Zurich, Switzerland, [38]University of Zurich, VP Medicine, Künstlergasse 15, 8001 Zurich, Switzerland
